# UnBlender: validating individual analyses in respiratory bulk RNA-seq cell type deconvolution

**DOI:** 10.64898/2026.06.01.727709

**Authors:** T. E. Gillett, M. van den Berge, M. C. Nawijn, G. H. Koppelman

## Abstract

Analysis of RNA-seq data of respiratory samples has contributed much to our understanding of lung disease. However, bulk RNA-seq data are dependent on both cell type composition and the transcriptional activity of these samples’ constituent cells, which complicates interpretation. Cell type deconvolution is frequently used to estimate cell type proportions of bulk transcriptomic gene expression data and improve interpretation of bulk transcriptomics data. However, accuracy of the estimated cell type proportions reported after deconvolution is unknown, which may have a negative impact on the validity of the conclusions drawn. Here, we present UnBlender, a pipeline that enables respiratory scientists to perform cell type deconvolution and routinely evaluate deconvolution accuracy of their approach. UnBlender allows for custom cell type deconvolution tailored to the research question at hand, using consensus cell type labels and validating the approach to promote accurate, reproducible results.

## Introduction

Lung tissue is complex: the lung consists of more than 50 annotated cell types (1,2). Many airway and lung diseases are the result of impaired cell function or dysregulated interactions between different cell types, including epithelial, mesenchymal and/or immune cells.

Analysis of RNA-seq data of airway and lung samples has contributed much to our understanding of respiratory disease (3). However, the bulk transcriptional profile of airway and lung tissue is dependent on both cell type composition and the transcriptional activity of the constituent cells. This complicates the interpretation of bulk RNA-seq data from these samples. Ignoring gene expression variation present in RNA-seq data due to differences in cell type abundance may hamper interpretation of changes in transcriptional activity in mixed tissues, and lead to false positive and false negative results (4). Whilst the still relatively new technology of single cell transcriptomics addresses this problem, bulk RNA-sequencing offers comprehensive assessment of tissue samples, is much less costly, and many datasets already exist. However, single cell reference data can be used to estimate cell type composition of a given bulk RNA-seq tissue sample through cell type deconvolution. This has been shown to be a useful approach to improve interpretation of bulk transcriptomics. Cell type deconvolution of bulk RNA-seq data into cell type proportions may constitute an outcome of an analysis (i.e. relative differences in cell type frequencies between health and disease), or can be used to improve analyses, e.g. when used as a covariate in differential gene expression or expression Quantitative Trait Locus analysis on the same bulk RNA-seq samples (5,6).

Cell type deconvolution uses a signature matrix to estimate the cell type proportions of the experimentally measured bulk transcriptomic gene expression data. These proportions can be estimated using a variety of methods, as described and reviewed previously (7-10). The signature matrix contains cell-type-specific gene expression values of a selection of genes that allow us to use bulk RNA-seq gene expression data to estimate the relative proportion of cell types in a sample. Suitable genes are abundantly and selectively expressed in a single cell type, but not or barely in others. Thus, the availability and accuracy of a matrix containing distinctive cell-type-specific gene expression data for the cell types in a bulk RNA-seq sample is crucial for the quality of cell type deconvolution.

Cell type deconvolution analysis of bulk RNA-seq faces several challenges, including the choice of cell types to be used for deconvolution and the lack of data needed to verify the accuracy of the signature matrix for the gene expression profiles of the selected sample type. The selection of cell types for deconvolution to be included in the signature matrix should ideally be motivated by known biology of the tissue sampled for bulk RNA-seq and the research question at hand but is also dependent on the availability of reference data. Cell types can be defined at multiple resolutions (e.g. “T-cells” versus the more specific but less frequent and transcriptionally related “CD8+ T-cells” and “CD4+ T-cells”). Recently, increasingly large and detailed reference datasets have become available, such as the Human Lung Cell Atlas (HLCA) and LungMAP Human Lung CellRef atlas (1,2), which improve the potential to extend the resolution of the signature matrix to less frequent cell types.

However, deconvolution of bulk transcriptomics into higher resolution cell types, especially when including multiple transcriptionally similar or related cell types, may lead to worse performance (11). Besides the lack of distinctive cell-type-specific genes for transcriptionally similar cell types, inclusion of low proportion cell types also affects cell type deconvolution accuracy, with the minimum percentage that is detectable varying considerably between deconvolution methods (11,12). Therefore, the accuracy of the estimated cell type proportions reported after deconvolution is unknown, with potentially far-reaching impact on the validity of the conclusions drawn. Accuracy of deconvolution algorithms and signature matrices can be validated using so-called gold standard datasets, which need to contain both bulk transcriptomics and cell type proportion measurements of the same samples. Lack of such gold standard datasets is an important challenge to cell type deconvolution of many bulk RNA-seq datasets (13,14).

As such, the question of which combinations of cell types can be deconvoluted accurately, at what cell type label resolution, within the context of a specific tissue and sample type, is usually left unaddressed. To this end, we have developed UnBlender, a pipeline that allows the respiratory community to routinely evaluate signature matrix deconvolution accuracy for data generated from bronchial biopsies, bronchial or nasal brushings and lung parenchyma resection samples. UnBlender offers a flexible framework to generate signature matrices based on consensus cell type labels across multiple resolutions, evaluates accuracy of their outcomes, and generates cell type proportion estimates in a given bulk-RNA-seq dataset of airway or lung tissue. The pipeline is currently available as a command line tool (https://github.com/Nawijn-Group-Bioinformatics/UnBlender) and will soon be accessible via a user-friendly graphical interface.

### Description of the Resource

#### Workflow

UnBlender generates deconvolution signature matrices based on consensus cell type labels of the Human Lung Cell Atlas across multiple resolutions in nasal, bronchial and lung parenchyma samples (1). The UnBlender workflow generates a custom signature matrix and evaluates its deconvolution accuracy in three steps (Figure 1). In step 1, the user is asked to select the sample type: bronchial biopsy, bronchial brush, nasal brush, or lung parenchyma resection. Based on the sample type, the user can select a combination of cell types of interest to deconvolute, at a chosen resolution. The pipeline then generates a custom signature matrix for this cell type selection, based on the HLCA scRNA-seq reference data (1). As specificity of genes used for deconvolution depends on the context of the sample and the other cells present in it, the reference scRNA-seq data from the HLCA are used to select the optimal set of cell-type-specific genes for the group of cell types selected by the user. UnBlender also generates HLCA-derived pseudobulk samples based on scRNA-seq data of the selected sample type, to resemble bulk RNA-seq samples for validation. During step 2, UnBlender deconvolutes these pseudobulk samples, of which the true cell type composition (the ground truth) is known, using the custom signature matrix. The command line interface implementation uses CIBERSORTx for signature matrix construction and deconvolution (15). The UnBlender pipeline reports on deconvolution accuracy by comparing the estimated cell type proportions to the known true proportions of the pseudobulk samples, providing measures of correlation and mean absolute proportional error (MAPE) for each of the selected cell types. This allows the user to evaluate whether their preferred selection of cell types is able to yield accurate results given the sample type used for bulk RNA-seq analysis. The output of step 2 of an example UnBlender run is provided in Supplementary Figure 1. If the deconvolution accuracy is sufficient, cell type deconvolution is performed on the bulk RNA-seq data of interest in step 3. If the deconvolution accuracy is insufficient, adjustments can be made to the cell type selection of step 1. More details on signature matrix construction, deconvolution, and evaluation are provided in the supplementary methods.

**Figure 1:**
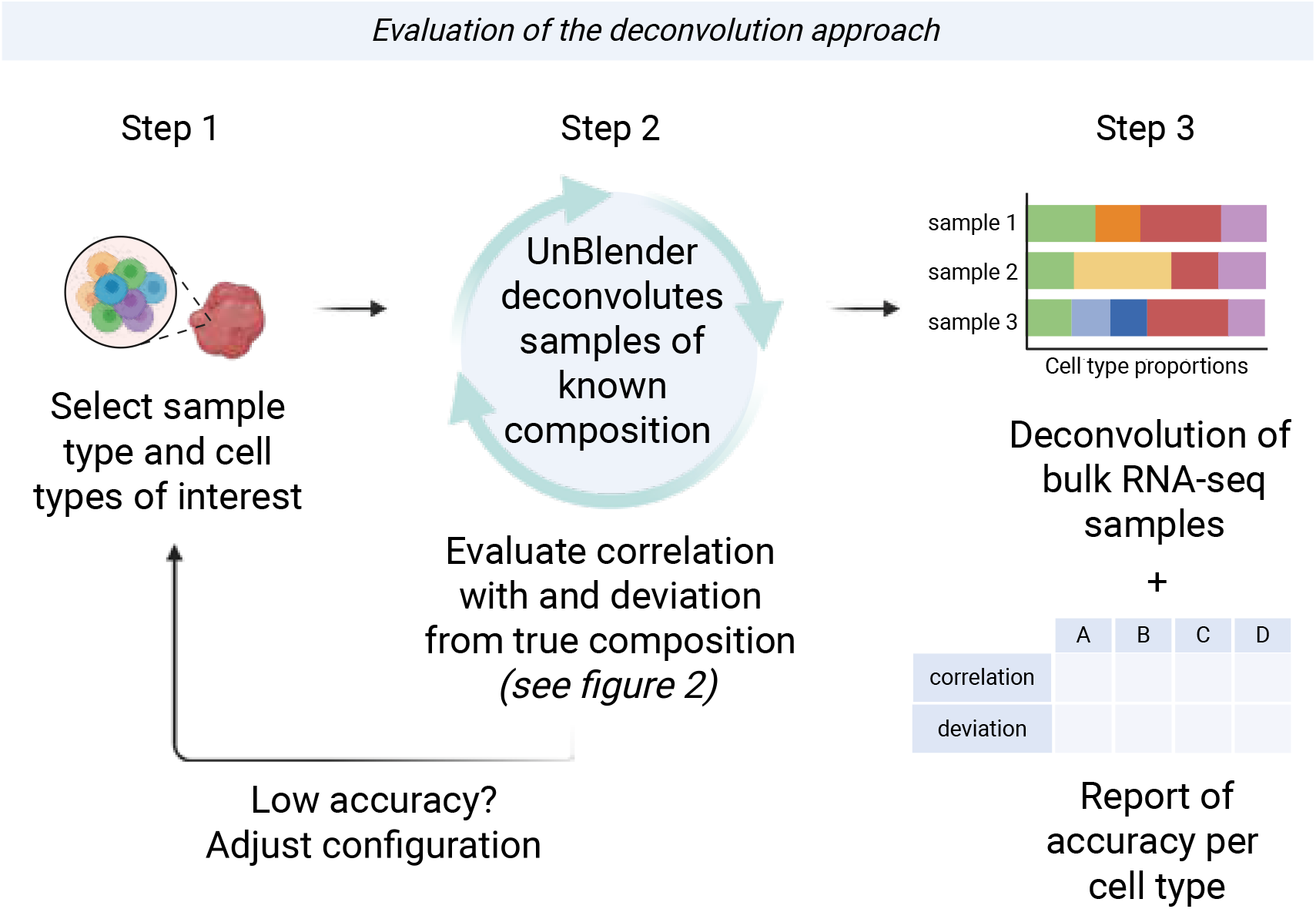
The UnBlender workflow evaluates whether a given cell type deconvolution approach is feasible: it deconvolutes pseudobulk samples of the relevant sample type and compares estimated proportions to the known pseudobulk composition. This allows the user to optimize their deconvolution approach to prevent inaccurate results before performing deconvolution on their own bulk RNA-seq data of interest.

We optimized the deconvolution workflow and established that cell type deconvolution accuracy stabilizes at the inclusion of 300 reference cells times the number of deconvoluted cell types (Supplementary Figure 2). Other optimization strategies were examined, including balanced sampling of the number of reference cells per cell type, and correction of deconvolution estimates for variation in cell-type-specific gene expression levels. Not all cell types present in a sample are necessarily included in each analysis, either due to low deconvolution accuracy or lack of relevance to the research question. Therefore, we also explored whether inclusion of other cell types, expected to be present as based on the sample type, improved deconvolution accuracy of the user selected cell types. Balanced sampling, correction for cell-type-specific expression and inclusion of additional cell types yielded no consistent improvements in deconvolution accuracy across cell types and were therefore not implemented in UnBlender (Supplementary Figure 3).

#### Output and evaluation

The UnBlender pipeline yields two measures of cell type deconvolution accuracy for each user-selected cell type: the Pearson correlation coefficient between the estimated cell type proportion and the known true cell type proportion, and the mean absolute proportional error (MAPE) of the estimate compared to the true proportion for that cell type (Supplementary Figure 2B). As shown in the case study using UnBlender to deconvolute nasal brush data in Figure 2 (and in similar analyses on bronchial brush, bronchial biopsy and parenchymal biopsy data, Supplementary Figure 4), deconvolution accuracy is variable between cell types. Therefore, deconvolution accuracy is evaluated for each cell type separately in UnBlender.

**Figure 2:**
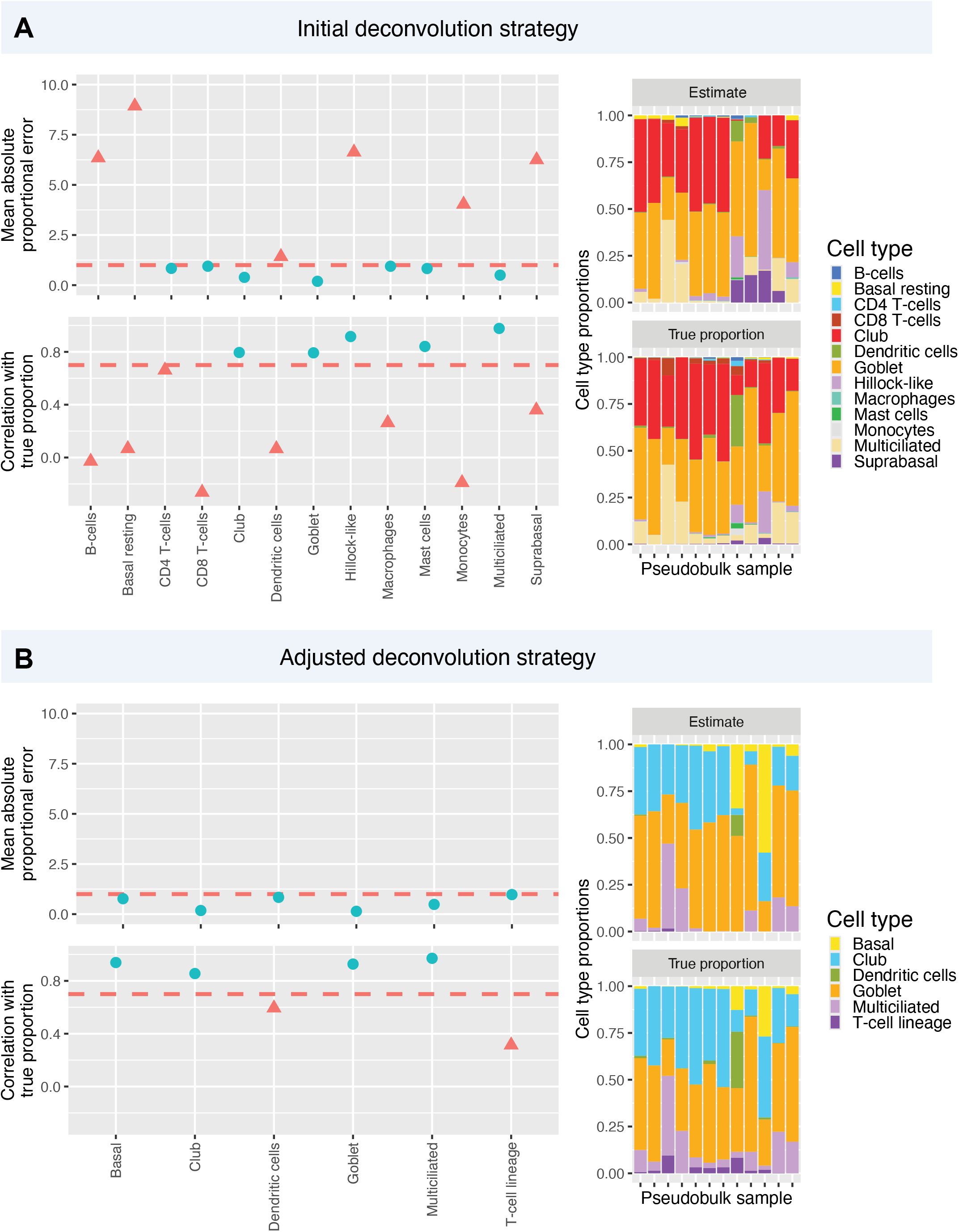
Case study: evaluating a deconvolution approach with the UnBlender pipeline, generated through deconvolution of 12 nasal brush pseudobulk samples. A) Pipeline output for an initial deconvolution strategy shows that deconvolution accuracy is highly variable between cell types. B) Deconvolution accuracy is improved after adjustment of the deconvolution strategy, reducing the granularity of the deconvolution matrix. For ease of interpretation the points have been separated into high versus low accuracy for correlation values above/below 0.75 and MAPE values of below/above 1.0, as indicated by the dashed lines.

Both MAPE and correlation are presented in the output, and are not necessarily in agreement. For example, in Figure 2A, the deconvolution of the CD8 T-cells is relatively accurate in terms of proportional error, but the correlation coefficient is below zero. In contrast, for the hillock-like cells a high correlation of estimated and true cell type proportions is observed, whilst the MAPE is relatively high. MAPE and correlation are two relatively easily interpretable metrics that represent separate aspects of how the estimated cell type proportions can be interpreted. Whilst MAPE measures the accuracy of the cell type proportion estimate in terms of its size, relative to the true cell type proportion, the correlation coefficient measures whether the estimates of multiple samples are an accurate indication of their relative differences. Whilst a suggestion is made as to which MAPE and correlation coefficient values may be considered ‘accurate’ and which warrant reconsideration of the deconvolution approach, the UnBlender interface allows the user to decide cut-offs for inclusion of cell types. UnBlender provides a table of accuracy metrics per cell type that can be reported in a scientific manuscript.

### Strategies to improve deconvolution accuracy

If the accuracy of the initial deconvolution approach is insufficient, the cell type selection used for the deconvolution needs to be adjusted and retested in an iterative fashion. First, it is useful to decide whether it is the MAPE or correlation scores that are most relevant to the intended downstream use of the deconvolution results, and how accurate MAPE or correlation scores need to be. This may depend on the research question, the sample size of the bulk RNA-seq experiment and the ability to verify the analysis results using independent methodology. Another consideration is whether an all-round high accuracy is required, or whether deconvolution accuracy of one or more specific cell types of interest can be optimized and that of the other cell types is less important.

For the initial analysis, users may want to select high resolution cell type labels for those cell types that are most relevant to the research question, e.g. AT2 cells rather than the more generic label of alveolar epithelial cells. However, it is important to realize that selecting low resolution cell type labels may yield more accurate results, as they constitute a larger proportion of the cells in a sample. Additionally, the chosen cell types must have distinct gene expression profiles that allow them to be distinguished from both similar and dissimilar cell types also selected for deconvolution. This is especially relevant for subsets of lymphoid, myeloid or secretory epithelial cells. When encountering poor performance of cell type deconvolution for such cell types, one approach is to merge two or more high resolution cell types with similar gene expression profiles into a single category (for instance, merge CD4 and CD8 T-cells into a generic T-cell label). Another approach is to remove those low proportion cell types that were not deconvoluted accurately from the analysis altogether. Figure 2B shows an improved cell type deconvolution strategy for the nasal brush samples, in which the resting basal, suprabasal and hillock-like cell type labels have been merged into the lower resolution ‘basal cell’ category. Additionally, cell types that constitute almost negligible proportions of nasal brush samples, resulting in low deconvolution accuracy, i.e. B-cells, macrophages, monocytes and mast cells, have been removed. As a result, MAPE has improved across tested cell types, indicating that deconvolution of any bulk RNA-seq data will yield more robust results when using the now validated cell type signature matrix.

## Discussion

UnBlender is a bioinformatic pipeline that allows users to generate a signature matrix for cell type deconvolution, evaluate the accuracy of its deconvolution outcomes as a routine step in the deconvolution workflow, and apply it to any bulk RNA-seq dataset generated from bronchial biopsies, bronchial brush, nasal brush and lung parenchyma resection samples. The tool enables validation of the deconvolution results by evaluating the accuracy of the deconvolution results each time a new cell type selection is used to generate a signature matrix for a given sample type.

The use of cell type deconvolution is becoming more widespread, as scRNA-seq data provide reference profiles of many respiratory cell types. We and others have observed variations in accuracy of the signature matrices used for deconvolution, which often remain unaddressed (14). The case studies presented here (Figure 2, Supplementary Figure 4) show the risk for low accuracy of deconvolution of respiratory bulk transcriptomics data into the constituent cell types, illustrating the need to validate the robustness of the preferred deconvolution strategy. It is difficult to predict a priori which cell types can be deconvoluted at which level of granularity. Deconvolution accuracy of a given cell type is not just influenced by the other cell types present in the sample, but also by which cell types are included in the analysis. Relevant factors include the distinctiveness of the cell type specific transcriptional profiles - a challenge that may be especially relevant in e.g. bronchial epithelial tissue with its many transitional cell states - and the cell type abundance within the sample type. Our findings in optimizing the deconvolution workflow indicate that the size of the reference data used to establish the signature matrix is less relevant for accuracy of deconvolution. Other challenges to accuracy of deconvolution include a potential lack of generalizability of healthy-reference-derived cell type signatures for samples obtained from individuals with lung disease or across donor demographics; and the need for a shared cell type nomenclature to establish a consensus on cell type identities within the respiratory field and thereby improve reproducibility between studies (13,14).

The differences in cell types’ total RNA content per cell, which often are disregarded during deconvolution, may also pose a challenge to deconvolution accuracy (13). Similarly, inclusion of lower numbers of reference cells in constructing the signature matrix may hamper the accurate deconvolution of cell types constituting a low proportion of the sample type. Finally, when not all cell types in a sample are included in the deconvolution analysis, part of that sample’s transcriptomic contents is not accounted for in the signature matrix. We examined optimization strategies to address these effects, but found that they yielded no consistent improvements in deconvolution accuracy across cell types.

UnBlender’s evaluation method is independent from both the deconvolution algorithm and reference dataset used, and can be adapted to improved deconvolution algorithms or reference datasets, such as a next iteration of the HLCA, as they become available. The deconvolution algorithm used in UnBlender was selected based on previous benchmarking efforts, which consistently showed good performance of ClBERSORTx (5,9).

The interpretation of the deconvolution accuracy and the selection of the cell types for the final deconvolution analysis is dependent on the context and research question of the study that is being performed. The metrics used for evaluation, correlation and MAPE, reflect distinct aspects of deconvolution accuracy. MAPE reflects the mean deviation of the deconvolution estimate relative to the cell type proportion, and is relevant when estimating the proportion of a cell type in a set of samples; whereas the correlation coefficient indicates whether the estimated cell type proportion across samples accurately reflects the relative compositional differences between the samples, and should be used when the goal of an analysis is to compare differences in cell type proportions between groups of samples. lmportantly, UnBlender does not suggest specific cut-offs for these accuracy metrics: what MAPE or correlation value is acceptable should be evaluated in the context of their downstream use, including research question, sample size, and whether analysis results may be independently verified. lmportantly, both correlation and MAPE are relatively easily interpretable, with a MAPE value of 0.5 indicating a deviation from the ground truth the size of half the ground truth proportion. These metrics can be reported in the method section of manuscripts using cell type deconvolution as part of their analysis, allowing for objective evaluation of the robustness of the presented results.

Finally, it is important to address a limitation of our approach. The MAPE and correlation metrics provided by UnBlender do not directly inform on the deconvolution accuracy of the bulk RNA-seq dataset in question. Factors such as donor demographics, ancestry, disease state, and technical variation are known to affect deconvolution results, and will differ between the pseudobulk samples used by UnBlender to assess the accuracy of deconvolution and the bulk RNA-seq data under study. In the future, this will be addressed by reference atlases’ inclusion of cells from participants of varying demographics and ancestry, as well as from different disease cohorts. In addition, whilst scRNA-seq data follows the same distributions as bulk transcriptomics (16), technical, cross-platform variation between scRNA-seq based pseudobulk samples and true bulk RNA-seq will further affect deconvolution outcomes. For example, current deconvolution matrices still lack (eosinophilic) granulocytes, due to RNA degradation during preparation of the single cell suspensions used for analysis. Therefore, UnBlender results are intended as an evaluation of the validity of the deconvolution approach, rather than a prediction of the precise accuracy of its results in a given bulk RNA-seq dataset.

Previous reviews have indicated the lack of ground truth datasets, as well as the lack of standardized cell type nomenclature as key challenges to performing accurate deconvolution analysis (13,14). UnBlender enables the wider respiratory community to evaluate the validity of their analysis approach without the need for an orthogonal ground truth measurement, the existence which would defeat the purpose of most deconvolution analyses, whilst making use of the consensus cell type labels established in the HLCA. Ongoing initiatives to coordinate harmonization of cell type labels across world-wide single cell consortia will further increase reproducibility of cell type deconvolution analysis and facilitate comparisons between studies.

In conclusion, the UnBlender pipeline enables respiratory scientists to perform cell type deconvolution in a flexible framework using consensus cell type labels across resolutions to suit their research question, generating a custom signature matrix, and evaluating the accuracy of its results. This approach allows us to perform reliable cell type deconvolution using the increased granularity that atlas-level reference datasets offer, balancing the need for high resolution deconvolution analyses with the reliability of their results, and allows for the accurate cell type deconvolution of many existing and new bulk transcriptomic datasets.

