## Supplementary Materials for "UnBlender: validating individual analyses in respiratory bulk RNA-seq cell type deconvolution"

#### Supplementary Materials and Methods

##### *The UnBlender pipeline*

***Selection of the reference data***

To run the UnBlender pipeline, the user first selects the relevant sample type: nasal brush, bronchial brush, bronchial biopsy, or parenchymal tissue resection. Then, the user selects the cell types that will be included in the signature matrix for deconvolution. Cell types can be selected according to the consensus cell type labels used in the HLCA, which are structured in 5 levels of increasing granularity (1). Parameterization of the command line implementation of the UnBlender pipeline is performed through a YAML file.

The reference data that will be used to construct the signature matrix is then derived from the HLCA by subsetting the atlas to the selected cell types, including only cells from samples matching the specified sample type. This subset is randomly downsampled to a total number of cells equalling *n* times the number of cell types included in the analysis. By default, n = 300, though in the command line implementation this number can be adjusted. If desired, the user may further specify any number of genes to be removed from the reference data, so that the expression levels of these genes will not impact the deconvolution analysis. This may e.g. be relevant to correct for batch effects known to be present in the bulk RNA-seq data that is to be deconvoluted.

***Generating the pseudobulk samples***

Next, up to 20 pseudobulk samples are generated based on the HLCA data. This is done by summing gene expression values per gene of the cells of up to 20 individual scRNA-seq samples of the selected sample type, in order to create a single bulk-like gene expression profile. For each of these samples, the composition according to the cell type labels specified in step 1 is calculated based on the single cell data. The exact number of generated pseudobulk samples varies depending on data availability (Table S1). Donor demographics for the four types of pseudobulk sample are given in table S2.

| *Sample type* | *Number of pseudobulks* | *Selection criteria* |
|---|---|---|
| Nasal brush | 12 | All samples of >2500 cells |
| Bronchial brush | 8 | All samples |
| Bronchial biopsy | 14 | All samples of >2500 cells |
| Parenchymal tissue resection | 20 | Randomly sampled from all samples containing >2500 cells, to limit computational complexity of the analysis. |

*Table S1: The number of pseudobulk samples generated from the HLCA scRNA-seq data, used to evaluate deconvolution accuracy.*

| | | Bronchial biopsy | Bronchial brush | Nasal brush | Parenchymal resection |
|---|---|---|---|---|---|
| n | | 14 | 8 | 12 | 20 |
| age, mean (sd) | | 43.5 (13.1) | 30.1 (4.2) | 37.6 (11.5) | 44.5 (16.0) |
| | *missing, n* | | 1 | | 3 |
| sex, n (%) | female | 4 (28.6) | 5 (62.5) | 6 (50.0) | 10 (50.0) |
| | male | 10 (71.4) | 3 (37.5) | 1 (8.3) | 6 (30.0) |
| ethnicity, n (%) | black | 1 (7.1) | 1 (12.5) | 1 (8.3) | 13 (65.0) |
| | white | 13 (92.9) | 7 (87.5) | 8 (66.7) | |
| | asian | | | 1 (8.3) | |
| | latino | | | 1 (8.3) | |
| | mixed | | | 1 (8.3) | |
| | *missing, n* | | | | 1 |
| smoking status, n (%) | active | | | | 5 (25.0) |
| | former | 4 (28.6) | 3 (37.5) | 1 (8.3) | 1 (5.0) |
| | never | 10 (71.4) | 5 (62.5) | 6 (50.0) | 13 (65.0) |
| | *missing, n* | | | 5 | 1 |
| BMI, mean (sd) | | 25.1 (3.6) | 22.1 (2.1) | 23.8 (3.7) | 34.6 (7.9) |
| | *missing, n* | | | 1 | 11 |

*Table S2: Donor demographics and clinical parameters for the pseudobulk samples used by UnBlender to evaluate deconvolution accuracy.*

***Deconvolution of pseudobulk samples and evaluation of accuracy***

The pseudobulk samples are deconvoluted using the selected reference data, yielding cell type proportion estimates per pseudobulk, using CIBERSORTx, an algorithm that approaches the deconvolution problem through support vector regression, a machine learning method (2). UnBlender runs CIBERSORTx with default settings and no cross-platform batch correction. CIBERSORTx additionally outputs the signature matrix that is generated to perform deconvolution, which can be used to deconvolute the user's bulk RNA-seq data.

Per cell type, the cell type proportion estimates are compared to the known true cell type proportions of the same pseudobulk samples, using two metrics. The spearman correlation over the samples of the estimated and true fraction is calculated; as well as the mean absolute proportional error (MAPE) of the estimated fraction compared to the true fraction: $\frac{1}{n} \cdot \sum_{i=1}^{n} \left| \frac{\hat{y}-y}{y} \right|$ . Both metrics, once for each cell type included in the deconvolution analysis, are provided to the user to determine whether the deconvolution accuracy is sufficient. The pipeline output also shows the composition of the pseudobulk samples used in the evaluation analysis and the underlying data on which the MAPE and correlation per cell type are calculated (Figure S1).

***Deconvolution of the bulk data***

If deconvolution accuracy is deemed sufficient, the specified deconvolution strategy can then be used to deconvolute the user's bulk RNA-seq samples. If not, the user can adjust their cell type selection for the deconvolution analysis to improve accuracy. The command line interface provides the signature matrix generated by CIBERSORTx as its output. This can be used to perform cell type deconvolution of the user's bulk RNA-seq data in an independent manner, to maximize flexibility, and can be re-used for future bulk RNA-seq datasets. Additionally, a table containing the deconvolution accuracy metrics per cell type is provided, which can be included in a manuscript.

#### *Case study*

We performed an example evaluation of a deconvolution strategy on 12 HLCA-derived nasal brush pseudobulk samples using the UnBlender pipeline. The deconvolution analysis included the following cell types: basal, suprabasal, club, goblet and hillock-like epithelial cells; B-cells, CD4 and CD8 T-cells, dendritic and mast cells, and macrophages, monocytes. To illustrate a second, improved deconvolution strategy, we subsequently performed another deconvolution on the same samples, including only basal, club, goblet, multiciliated, dendritic and T-cells.
Similarly, we performed example UnBlender pipeline runs for 8 bronchial brush, 14 bronchial biopsy and 20 parenchymal biopsy pseudobulk samples. For the bronchial brush pseudobulk sample deconvolution, the following cell types were included: basal cells, secretory cells, multiciliated lineage cells, ionocytes, tuft cells, B-cell lineage cells, dendritic cells, mast cells, monocytes, alveolar macrophages, interstitial macrophages, and ILC/NK cells and T-cell lineage (combined). For the bronchial biopsy pseudobulk sample deconvolution, the following cell types were included: capillary endothelial cells, venous endothelial cells, B-cells, NK cells, CD4 T-cells, CD8 T-cells, dendritic cells, alveolar macrophages, interstitial macrophages, mast cells, monocytes, fibroblasts, smooth muscle cells, basal and secretory cells (combined), and multiciliated cells. For the parenchymal biopsy pseudobulk sample deconvolution, the following cell types were included: capillary endothelial cells, venous endothelial cells, arterial endothelial cells, lymphatic endothelial cells, AT2 cells, T-cell lineage, B-cells, dendritic cells, alveolar macrophages, interstitial macrophages, mast cells, monocytes, fibroblasts, smooth muscle cells, basal and secretory cells (combined), and multiciliated cells.

#### *Optimization of the UnBlender pipeline*

The following optimization strategies were tested, but ultimately not implemented in the UnBlender pipeline.

***Effect of the number of reference cells***

One important parameter to determine is the size of the reference data needed for accurate deconvolution. We therefore assessed the effect of the number of cells included in the reference data on the subsequent deconvolution accuracy. Nasal brush and parenchymal biopsy samples were selected to represent a relatively simple and more complex cell type composition scenario, respectively. The UnBlender command line implementation was run to evaluate deconvolution accuracy per cell type, selecting the following cell type labels to be included in the analysis. Nasal brush: basal resting and suprabasal cells (combined), club cells, dendritic cells, goblet cells, multiciliated lineage cells, and T-cell lineage cells; parenchymal resection: alveolar macrophages, AT2 cells, B-cell lineage cells, basal and secretory cells (combined), dendritic cells, arterial, capillary and venous endothelial cells, fibroblast lineage cells, ILC/NK and T-cell lineage cells (combined), interstitial macrophages, endothelial lymphatic cells, mast cells, monocytes, multiciliated lineage cells, and smooth muscle cells.

We ran this analysis by including a variable number of cells corresponding to the number of cell types times *n* total number of cells in the scRNA-seq reference data used to generate the signature matrix, for *n* between 50 and 1250, increasing in steps of 50. For the parenchymal biopsy samples, where 16 distinct cell types were included in the analysis, the CIBERSORTx algorithm was unable to run past *n* = 600. 25 replicate analyses were performed per value of *n* for each sample type.

***Balanced sampling of cell types***

To assess whether deconvolution of lower proportion cell types could be improved by increasing the number of reference cells used to generate the signature matrix, we reran the same analyses as described in the previous paragraph, but now sampled an equal number of 300 reference cells per cell type. Where fewer than 300 cells were available for a cell type, the total number of cells of that cell type available in the HLCA was included in the reference data. These results were compared to deconvolution accuracy results when randomly sampling a total number of cells equalling 300 times the number of cell types: 300*6 = 1800 for nasal brush samples; and 300*16 = 4800 cells for parenchymal biopsy samples. 25 replicate UnBlender runs were performed per analysis.

***Filler cell types***

Often a deconvolution analysis does not need to include the exhaustive list of cell types present in a sample in its reference data, either because certain cell types are not relevant to the research question, because of low cell type proportions, or when evaluating cell type deconvolution accuracy indicates the cell type can not be deconvoluted well. We explored whether inclusion of such cell types in the reference data used to generate a signature matrix improves the deconvolution accuracy of the cell types originally included in the analysis. For this purpose, B-cell lineage cells, fibroblasts, hillock-like cells, mast cells, monocytes and macrophages (combined), and rare epithelial cells were added to the previously described list of cell types for the deconvolution of nasal brush samples; for the parenchymal tissue deconvolution the following cell types were added: AT1 cells, and mesothelial cells. 25 replicate UnBlender runs were performed per analysis.

***Correction for cell type-specific expression levels***

To assess the effect of the varying average number of counts per cell type included in the reference data, we performed a correction for these cell type-specific expression levels. Having estimated cell type proportions for nasal brush samples and parenchymal biopsy samples as described under "*effect of the number of reference cells"* for the analysis including 300 reference cells per cell type, we then corrected these estimates by dividing the percentage by the mean number of counts per reference cell for that cell type, and then scaling the sum of the estimates back to 100%. This was done in two ways, either using the total counts over all genes included in the reference data, or including only those genes that were included in the signature matrix. 25 replicate UnBlender runs were performed per analysis.

### Supplementary Results

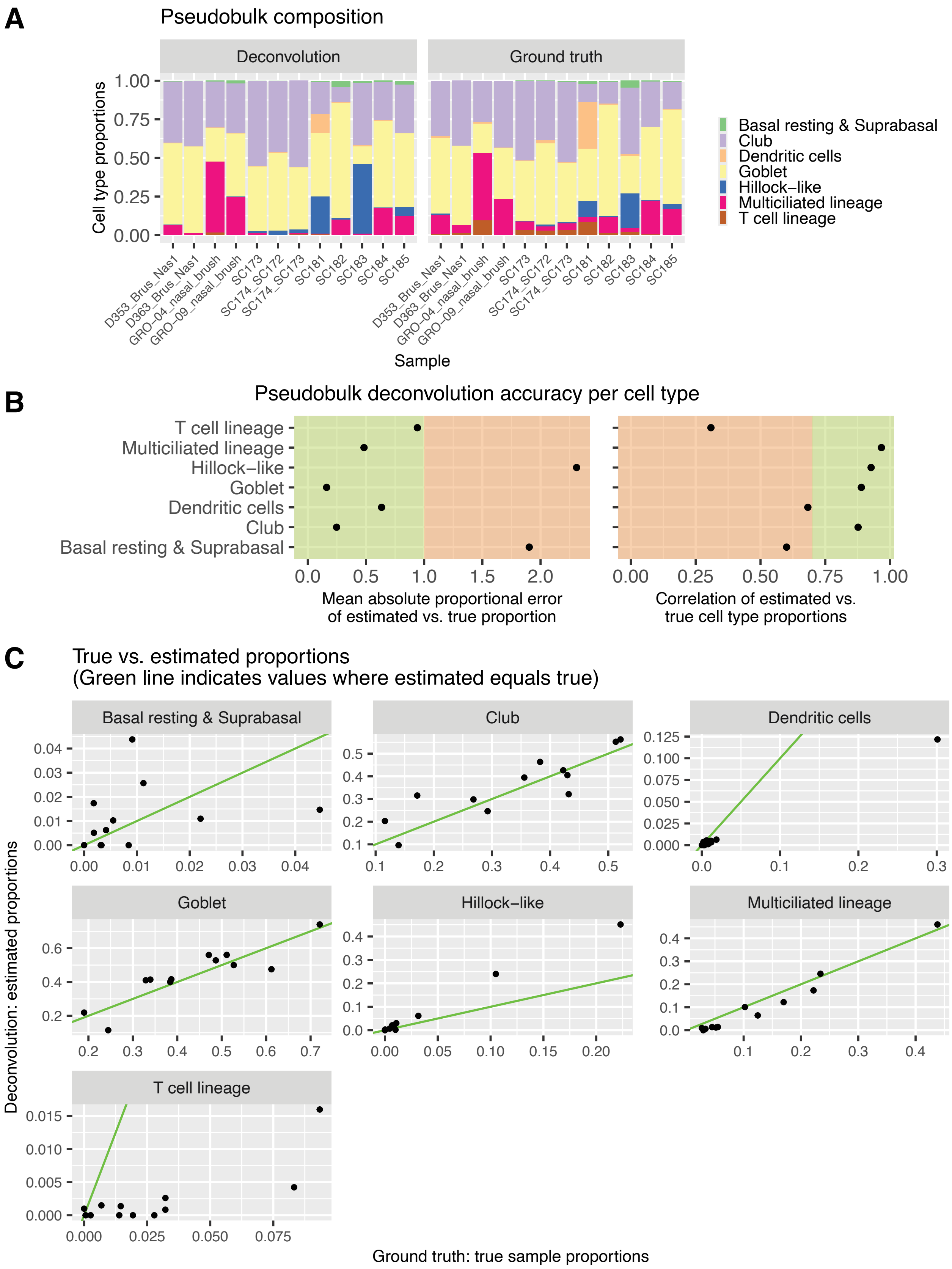

*Figure S1 (previous page): An example of the output provided by UnBlender to be used to evaluate deconvolution accuracy per cell type: pseudobulk cell type composition and estimated composition (A); MAPE and correlation of the true and estimated proportions per cell type; and (B) the underlying proportions per sample. Example analysis on 12 nasal brush pseudobulk samples, deconvoluting into 7 cell types.*

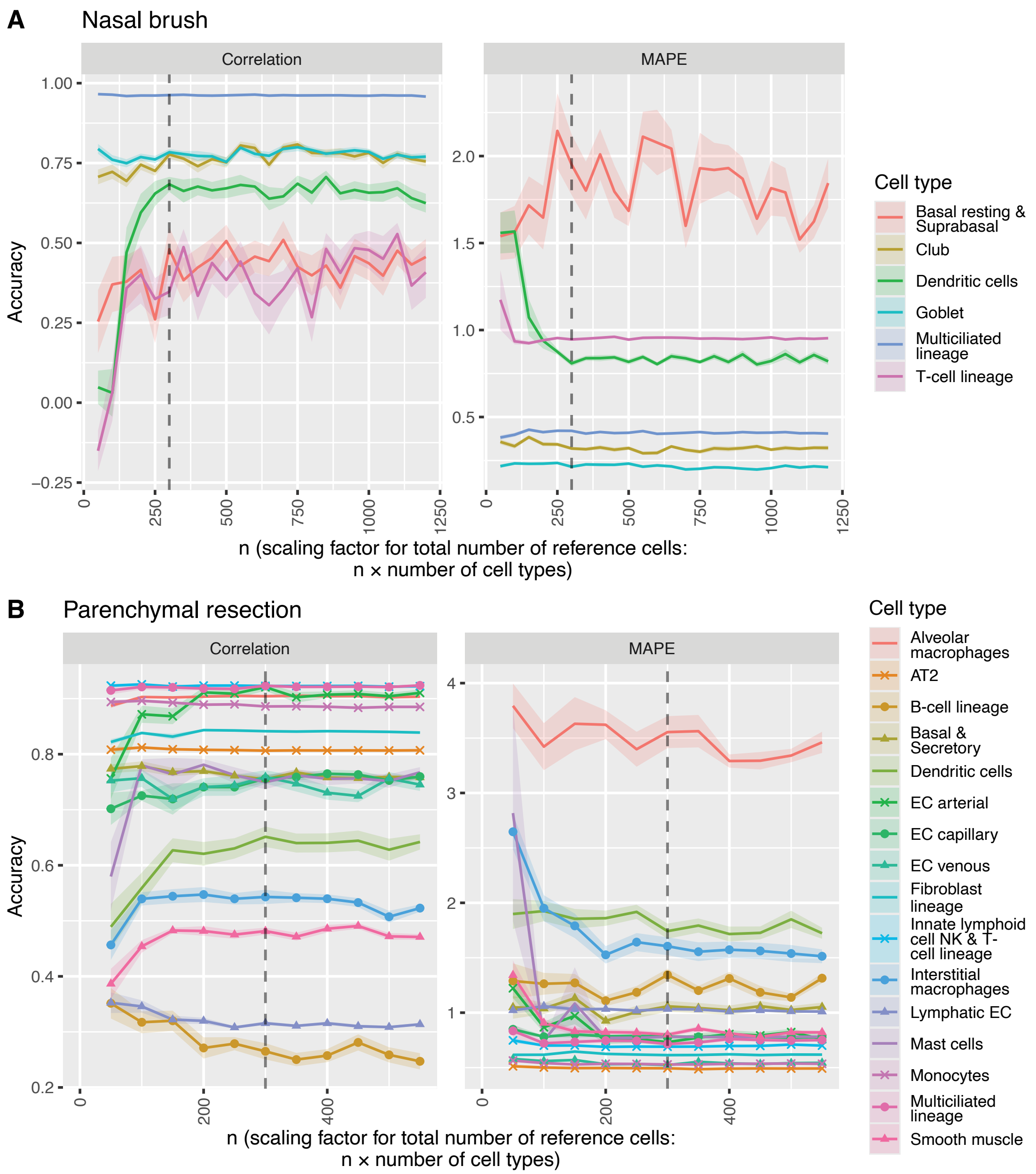


*Figure S2: Deconvolution accuracies after deconvolution of 12 nasal brush pseudobulk samples (A) and 20 parenchymal biopsy pseudobulk samples (B) over a range of varying numbers of cells included in the reference. The x-axis shows* n, *where* n *times the number of cell types is the total number of reference cells included. Accuracy is measured as both the MAPE and the correlation coefficient of the cell type proportion estimates compared to the known cell type proportions. Each colour represents a cell type included in the analysis.*

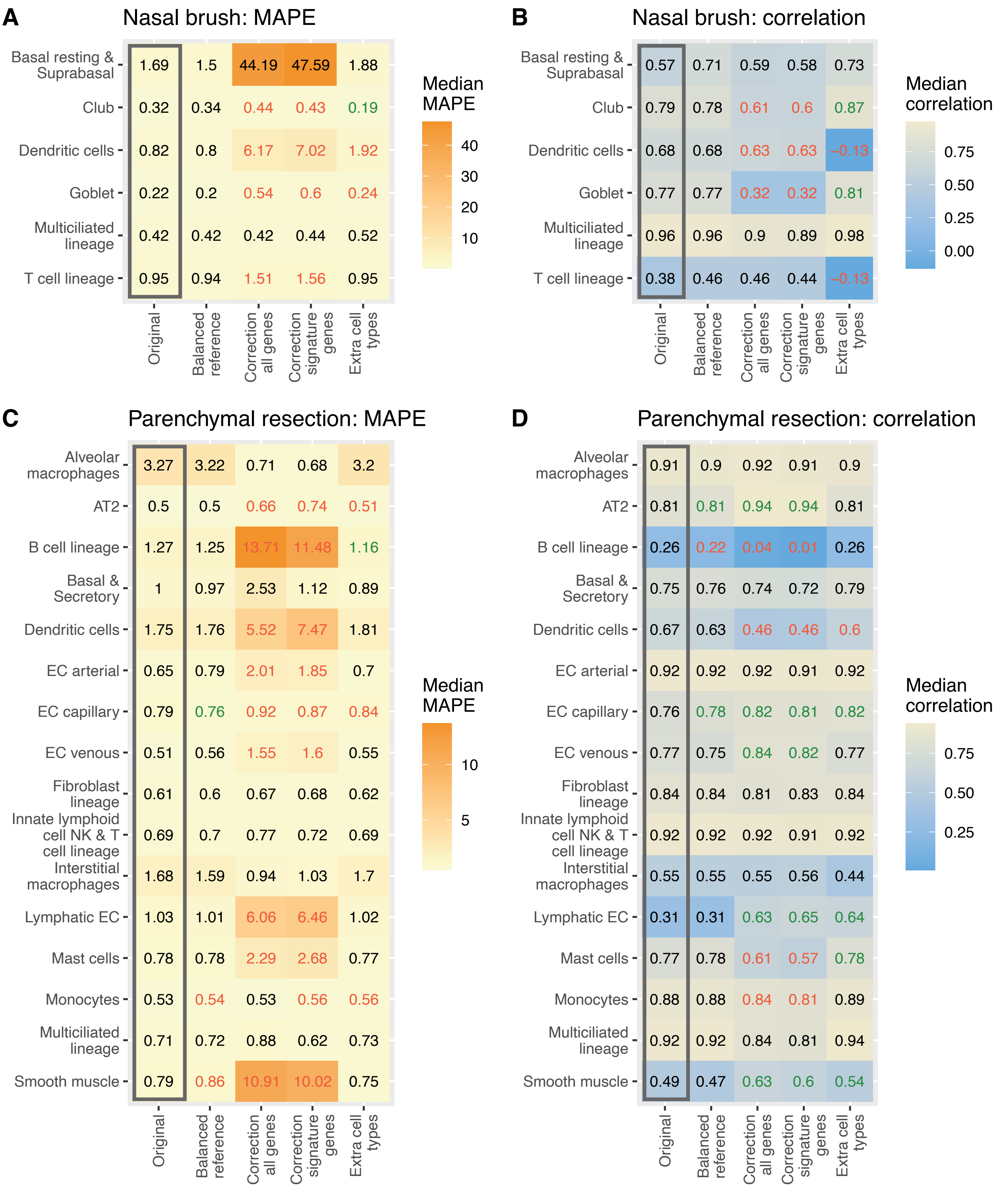
A
Nasal brush: MAPE
Basal resting & Suprabasal
Club
Dendritic cells
Goblet
Multiciliated lineage
T cell lineage
1.69 1.5 44.19 47.59 1.88
0.32 0.34 0.44 0.43 0.19
0.82 0.8 6.17 7.02 1.92
0.22 0.2 0.54 0.6 0.24
0.42 0.42 0.42 0.44 0.52
0.95 0.94 1.51 1.56 0.95
Original
Balanced reference
Correction all genes
Correction signature genes
Extra cell types
Median MAPE
40
30
20
10
B
Nasal brush: correlation
0.57 0.71 0.59 0.58 0.73
0.79 0.78 0.61 0.6 0.87
0.68 0.68 0.63 0.63 −0.13
0.77 0.77 0.32 0.32 0.81
0.96 0.96 0.9 0.89 0.98
0.38 0.46 0.46 0.44 −0.13
Median correlation
0.75
0.50
0.25
0.00
C
Parenchymal resection: MAPE
Alveolar macrophages
AT2
B cell lineage
Basal & Secretory
Dendritic cells
EC arterial
EC capillary
EC venous
Fibroblast lineage
Innate lymphoid cell NK & T cell lineage
Interstitial macrophages
Lymphatic EC
Mast cells
Monocytes
Multiciliated lineage
Smooth muscle
3.27 3.22 0.71 0.68 3.2
0.5 0.5 0.66 0.74 0.51
1.27 1.25 13.71 11.48 1.16
1 0.97 2.53 1.12 0.89
1.75 1.76 5.52 7.47 1.81
0.65 0.79 2.01 1.85 0.7
0.79 0.76 0.92 0.87 0.84
0.51 0.56 1.55 1.6 0.55
0.61 0.6 0.67 0.68 0.62
0.69 0.7 0.77 0.72 0.69
1.68 1.59 0.94 1.03 1.7
1.03 1.01 6.06 6.46 1.02
0.78 0.78 2.29 2.68 0.77
0.53 0.54 0.53 0.56 0.56
0.71 0.72 0.88 0.62 0.73
0.79 0.86 10.91 10.02 0.75
Median MAPE
10
5
D
Parenchymal resection: correlation
0.91 0.9 0.92 0.91 0.9
0.81 0.81 0.94 0.94 0.81
0.26 0.22 0.04 0.01 0.26
0.75 0.76 0.74 0.72 0.79
0.67 0.63 0.46 0.46 0.6
0.92 0.92 0.92 0.91 0.92
0.76 0.78 0.82 0.81 0.82
0.77 0.75 0.84 0.82 0.77
0.84 0.84 0.81 0.83 0.84
0.92 0.92 0.92 0.91 0.92
0.55 0.55 0.55 0.56 0.44
0.31 0.31 0.63 0.65 0.64
0.77 0.78 0.61 0.57 0.78
0.88 0.88 0.84 0.81 0.89
0.92 0.92 0.84 0.81 0.94
0.49 0.47 0.63 0.6 0.54
Median correlation
0.75
0.50
0.25

*Figure S3 (previous page): Cell type proportion estimate accuracy after deconvolution of 12 nasal brush pseudobulk samples (A, B) and 20 parenchymal biopsy pseudobulk samples (C, D) following different optimization strategies: sampling equal numbers of reference cells per cell type; correcting for cell type-specific expression levels of all genes, or of the genes included in the deconvolution signature matrix; and the inclusion of additional cell types in the analysis. Deconvolution accuracy without the optimization strategies ("original") is shown for comparison and marked by a grey box. Median correlation or MAPE values in green and red are significantly improved or worsened, respectively, compared to the corresponding "original" value.*

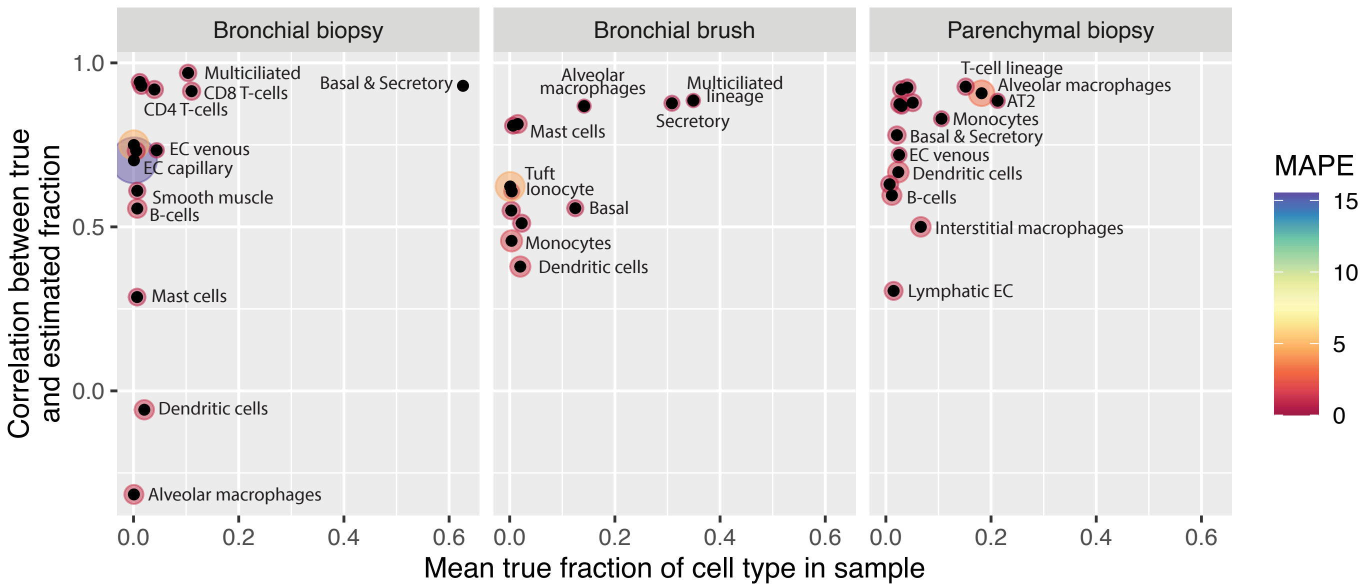


*Figure S4: Example deconvolutions of bronchial brush, bronchial biopsy and parenchymal biopsy samples show varying deconvolution accuracy per cell type. Each dot represents a cell type included in the analysis, with the correlation between the true and estimated cell type percentages on the y-axis, and the mean absolute proportional error (MAPE) size as indicated by the size and colour of the circle surrounding each dot.*
